# Label-free visualization of type IV pili dynamics by interferometric scattering microscopy

**DOI:** 10.1101/298562

**Authors:** Lorenzo Tala, Adam Fineberg, Philipp Kukura, Alexandre Persat

## Abstract

Type IV pili (TFP) are slender objects that assemble by polymerization and secretion of protein subunits from bacterial cell surfaces. The mechanisms by which these surface structures of microscopic length and molecular diameter modulate the physical interaction of bacteria with their environment, however, remains poorly understood largely due to limitations in our ability to monitor and characterize the dynamics of individual TFP. Here, we demonstrate that interferometric scattering microscopy (iSCAT) enables label-free and dynamic visualization of TFP in intact cells of the opportunistic pathogen *Pseudomonas aeruginosa*. As a result, we can directly monitor extension, attachment and retraction events on millisecond timescale and nanometer length scale in three dimensions. These capabilities allow us to observe that *P. aeruginosa* is able to crawl against the direction of flow using short TFP filaments. Also, careful observation show that TFP retract rapidly after surface attachment, suggesting that *P. aeruginosa* senses contact of the pilus tip with the solid substrate. These results illustrate the power of iSCAT for the label-free visualization of small, dynamic microbial extracellular structures.

## Introduction

The surface of bacteria is decorated with a variety structures, many of which allow mechanical interaction with the environment or with other cells^1,2^. Flagella and pili are long and thin filaments that extend from single cells by polymerization of elementary subunits. In contrast to type IV pili (TFP), a variety of microscopy methods have revealed the dynamics of the 20 nm-wide flagellum, for example by non-specific labeling with synthetic fluorescent dye^3^, or by label-free methodologies such as darkfield microscopy^4^.

TFP are long and thin filaments that extend from the poles of many bacteria^5^. In *Pseudomonas aeruginosa*, successive rounds of TFP extension, attachment and retraction power cell displacements known as twitching. This mechanical activity also activates surface mechanosensing, which promotes the production of multiple virulence factors^6^. TFP extend through rapid periplasmic polymerization of the major pilin subunit PilA by the extension motor PilB. Stepwise helical assembly translocates the filament through an outer membrane pore. The retraction motors PilT and PilU power PilA depolymerization by rapidly reinserting monomers into the inner membrane^5^. The mechanism by which TFP activity is regulated on short timescales of extension-retraction (subsecond) and on longer timescales of surface colonization and mechanically-activated transcription (minutes to hours), however, cannot be tested, due to a lack of suitable visualization techniques. Understanding how *P. aeruginosa* dynamically regulates TFP activity thus rests on the availability of methodologies capable of accessing both short and long timescales, without affecting cell viability.

TFP are clearly visible on electron micrographs of fixed cells which, however, are intrinsically static^7^. Indirect force spectroscopy methods such as optical tweezers, nanopillars and atomic force microscopy can provide dynamic measurements of the forces generated by TFP, which range between 10 to 100 pN^8–10^. Generation of fluorescent fusions to pilin subunits for direct visualization is complicated by the spatial requirement for extracellular translocation. A non-specific labeling method has successfully highlighted the dynamics of single extension and retraction events^11^. This technique, however, subjects cells to hour-long labeling and damaging washing procedures that affect the integrity of the fibers, does not provide long-term dynamics and may impair TFP dynamics by obstructing translocation. Moreover, since visualization relies on recycling of labeled-subunits, the signal decays over time from photobleaching and subunit turnover. Thus, measurements of TFP dynamics have been limited, resulting in many open questions concerning the regulation of TFP length, number, extension-retraction frequencies, orchestration and mechanosignaling.

TFP filaments are extracellular objects and do not interact with light sufficiently strongly to be observed with traditional label-free microscopy methods. We were enthusiastic when recent progress in interferometric scattering microscopy (iSCAT) has pushed the sensitivity of optical detection so far as to detect single, 6 nm-wide unlabeled actin filaments assembled *in vitro*^12^. iSCAT has also revealed the *in vitro* dynamics of microtubules^13^, myosin motors^14^, vesicle fusion^15^, and single virus translocation^16^ at high temporal and spatial resolutions by leveraging the interference generated by a reference light source with the light scattered by the sample. Visualization of the interferometric component increases the sensitivity to objects that scatter poorly, such as single proteins. In practice, the light reflected by the coverslip is used as a reference light source. The light scattered by the sample returns through the objective, thus interfering with the reflected light. In contrast to darkfield microscopy, the power of iSCAT resides in its increased sensitivity to small scatterers^17^. Here, we leverage the sensitivity of iSCAT to perform *in vivo*, label-free real-time visualization of *P. areuginosa* extracellular filaments, thereby solving a major limitation in our ability to characterize their structure and dynamics.

## Results

### iSCAT reveals bacterial extracellular structures

To demonstrate the capability of iSCAT to resolve small bacterial extracellular machinery, we first visualized *P. aeruginosa* cells and selected mutants in flagella and type IV pili components. Wild type (WT) *P. aeruginosa* under brightfield illumination appears as rod-shaped without any particularly visible structures (**Fig. 1A**). In contrast, simultaneous iSCAT imaging of the same cell reveals micrometer long extracellular filaments and shorter structures with alternating contrast extending from the cell body (**Fig. 1B**). iSCAT images of in-frame deletion mutants in the major pilin subunit gene *pilA* display none of these slender structures as shown in **Fig. 1C**, thereby demonstrating that they correspond to TFP. A deletion mutant in the flagellin gene *fliC* (**Fig. 1D**) did not display structures with periodic pattern, demonstrating that the helical shape of a single flagellum filament generates this signal. Measuring the wavelength of these patterns gives an estimate of 1.4 µm ± 0.1 µm (*N* = 10) for the helical pitch of the flagellum, consistent with previous measurements^18^. Since the signal generated by the flagellum is strong compared to TFP, we performed all subsequent visualizations in a *fliC*^-^ background.

**Figure 1.**
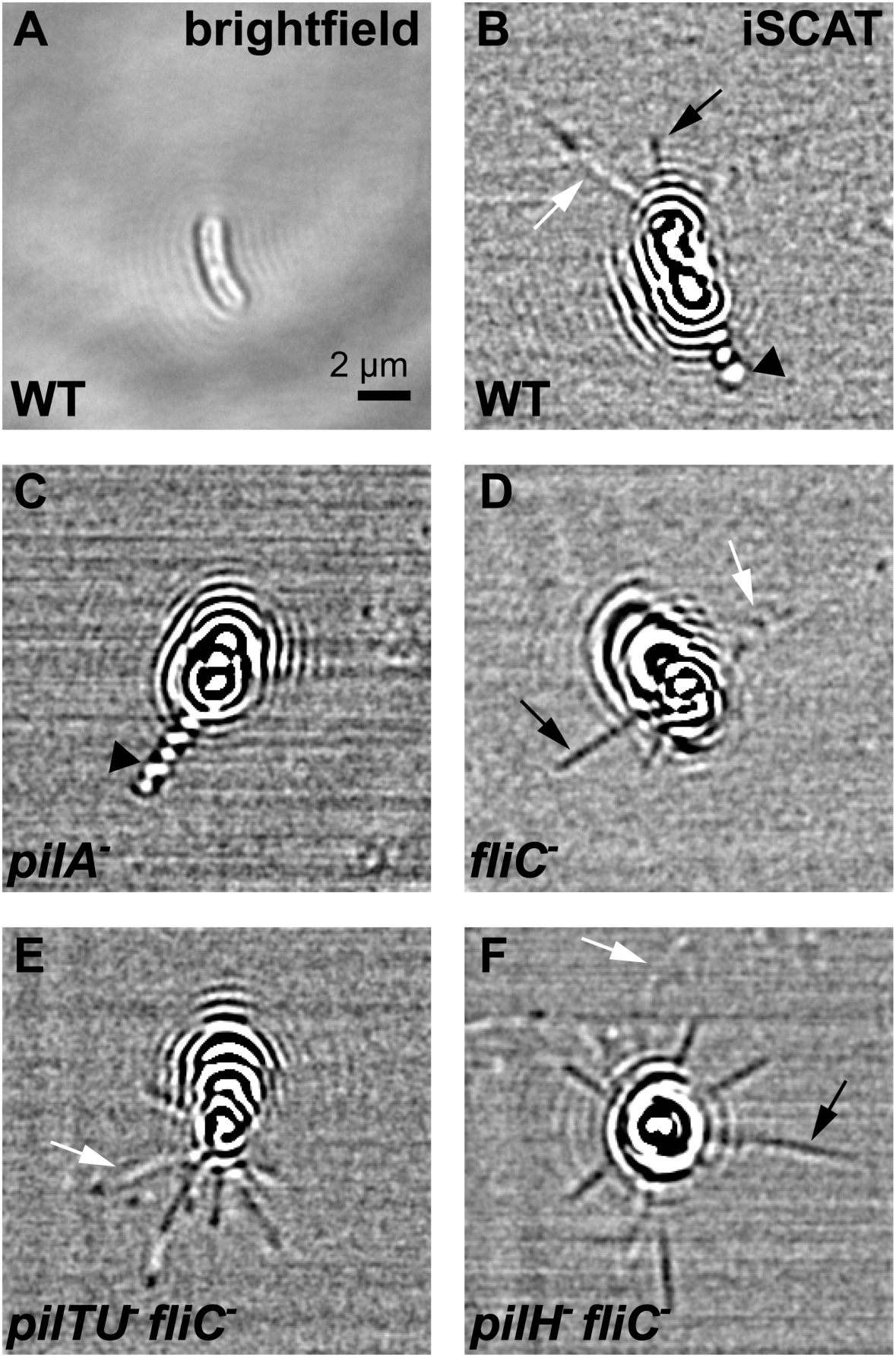
iSCAT reveals extracellular *P. aerignosa* filaments. (A) Brightfield and (B) iSCAT images of a single wild-type *P. aeruginosa* cell. The brightfield image only shows the rod-shaped body of the bacterium, but multiple extracellular structures appear in the iSCAT channel. This includes a polar structure with alternating dark and white contrast (black arrowhead) and slender structures with constant (black arrow) or spatially varying contrast (white arrow). (C) A deletion mutant of the major pilin subunit gene *pilA* displayed no extracellular slender structures, while (D) a mutant in the flagellin subunit gene *fliC* showed no polar filament with alternating contrast. These two classes of signals can thus be respectively attributed to TFP and flagella. (E) A strain lacking retraction motor genes *pilT* and *pilU* has many TFP. (F) A strain lacking the gene coding for the Chp response regulator *pilH* also displayed many TFP but with generally constant negative contrast.

To further demonstrate the power of iSCAT in visualizing TFP, we studied two mutants that display increased levels of surface TFP^19^. First, we imaged a deletion mutant in the retraction motors genes *pilT* and *pilU* which are able to extend, but not retract TFP. We observed many TFP in this strain, generally 5 to 10, extending from each cell pole (**Fig. 1E**). We separately imaged a deletion mutant in the response regulator gene *pilH*, which displays increased single cell motility compared to WT^20^. Consistent with this, our visualizations confirm that *pilH*^-^ mutant has many extended TFP (**Fig. 1F**).

In all strains, we observed three distinct patterns generated by TFP. They either appeared as straight dark filaments, straight filaments with alternating black and white contrast, or curved filaments of low intensity (**Fig. 2**). Changes in contrast of a fiber are a manifestation of the shift between constructive and destructive interference with a full phase change corresponding to a spatial shift of λ/2*n* where λ is the illumination wavelength and *n* the index of refraction of the medium^17^. Thus, contrast values can be used as a proxy for in depth position of TFP^21^. TFP laying flat against the surface have uniform intensity (**Fig. 2A**). The ones that are at an angle exhibit changes in contrast with successive positive and negative intensity values (**Fig 2B and C**). For example, the three successive contrast changes in **Fig. 2B** indicate that the straight filament extends approximately 360 nm above the glass surface, likely originating from the side of the cell pole. In our visualizations, we noticed that in absence of retraction motors, TFP appear as curved filaments with changes in intensity (**Fig. 1E**), while TFP from strains that can retract TFP most frequently appear as dark filaments (**Fig. 1B-D-F**). This suggests that retraction orients TFP parallel to the glass surface promoting adhesion along their length rather than uniquely at the tip. Consistent with this, TFP of *pilTU*^-^ *fliC*^-^ attach from their tips but remain floppy, never transitioning to the tensed state (**Fig. 1E**, **Movie 1**).

**Figure 2:**
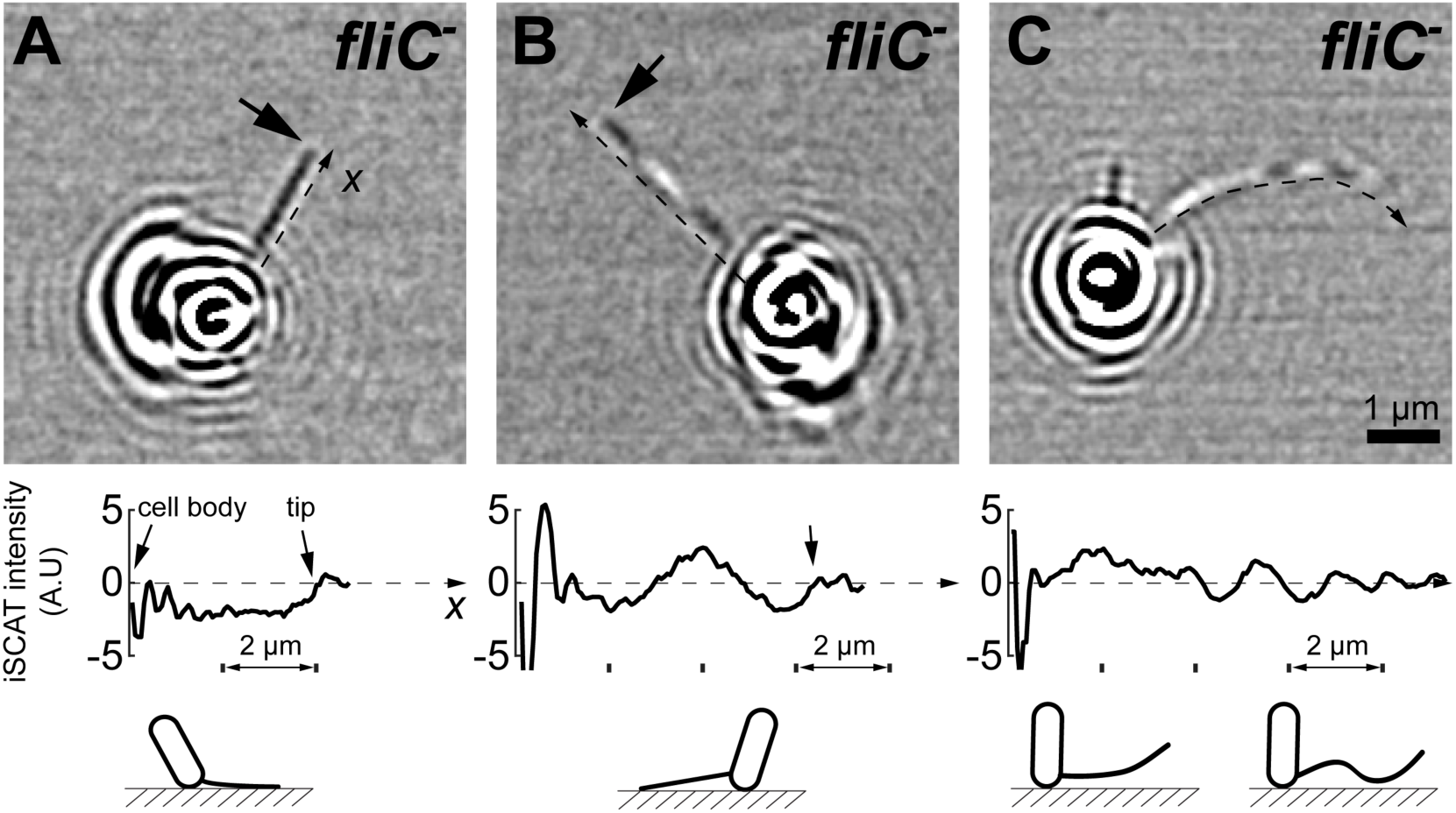
Inferring TFP orientation from iSCAT signal intensity. The signal generated by iSCAT through constructive or destructive interference appears as a change in contrast from the background, which depends on the position of the scattering object. (A) TFP oriented parallel to, and against the glass surface appear as a dark line. The intensity along the fiber starting from cell body to tip has approximately constant negative values. The signal spikes at the vicinity of the cell are generated by the strong scattering of its body. (B) Straight TFP appearing with alternating signal polarity indicate that the fiber extends away from the surface. The contrast values show three polarity changes (dark to clear), which corresponds to a total height change of 360 nm in the direction perpendicular to the imaging plane. We thus estimate that the pilus likely extends from the side of the cell pole. (C) Many TFP appear with large spatial and temporal fluctuations in iSCAT signal, indicating that they remain unattached. Their spatial orientation cannot be strictly inferred directly from iSCAT intensity as the signal can be generated from multiple configurations as illustrated under the intensity graph.

### TFP activity

Multiple pathways regulate the production of TFP components^5^. A clear understanding of how these modulate TFP length, orientation, extension-retraction frequency and attachment, however, is lacking. Our understanding of TFP activity regulation by environmental signals, and consequently of its function in an ecologically-relevant context, is therefore limited. To demonstrate the capability of iSCAT to reveal biophysical components of TFP dynamics, we optimized iSCAT to reduce phototoxicity by using higher iSCAT and autofocus laser wavelengths and a mechanical shutter, thereby enabling visualization of single cells over several minutes of surface exploration ^22^. These improvements allowed us to capture multiple successive extension-retraction events while monitoring cell body displacements. **Movie 2** shows a 10 min visualization of the hyperactive TFP mutant *pilH*^-^ *fliC*^-^ at 2.5 frames per second exhibiting frequent TFP attachment and retraction events. We isolated each cell within this sequence and counted attachment-retraction events to determine their associated frequencies (**Fig. 3**). For the first and second cell, we counted 15 retractions within 2 min (**Fig. 3A**) and 29 events in 3 min (**Fig. 3B**), respectively, yielding retraction frequencies of 7 to 10 per minute. This sequence also shows that attached TFP can either remain under tension without retracting, or retract under tension by sliding onto the surface.

**Figure 3:**
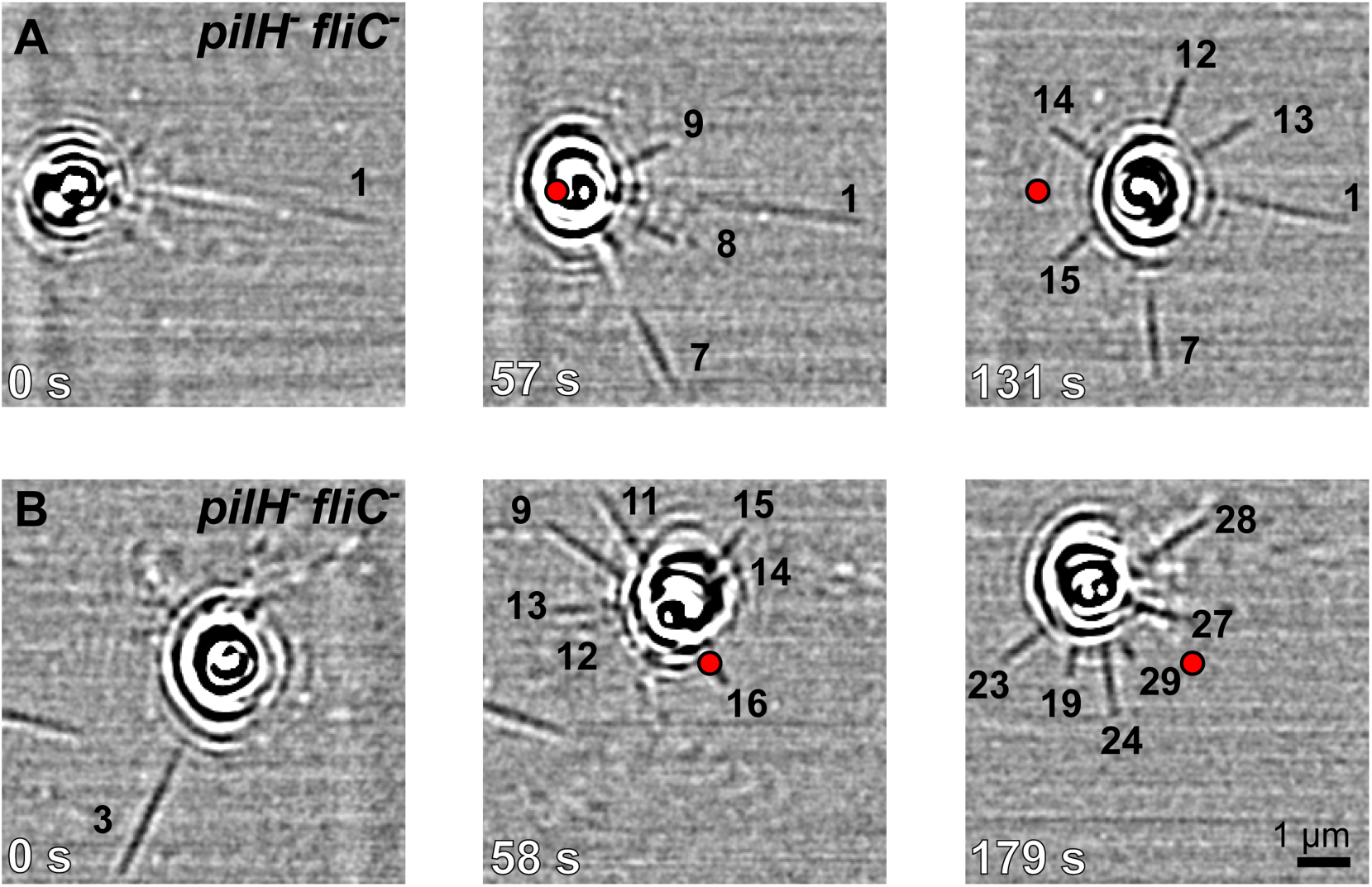
Tracking long-term TFP activity of single cells with iSCAT. We visualized TFP activity of a *pilH^-^ fliC^-^* mutant with constitutively high surface motility phenotype. In these visualizations, we can perform tracking of TFP attachment and retraction while monitoring corresponding cell trajectories. In (A), we measured 15 attachment-retraction events during the 2 min visualization, corresponding to 7 events per minutes. We tracked and numbered some of the events on the image sequence. The red dot indicates the initial position of the cell body. We performed a similar analysis on the cell shown in (B) and measured an attachment-retraction frequency of 10 events per minute. **Movie 2** is a dynamic visualization of the same data.

### Upstream migration

*P. aeruginosa* migrates upstream in flow environments in a TFP-dependent manner^23^. How TFP power this migration remains unclear, however, as intuitively TFP should orient in the direction of the flow before attaching to the surface. Visualizing extension and retraction events under flow within a microfluidic channel confirmed TFP power upstream twitching as some filaments extended against the direction of flow (**Fig. 4, Movie 3, 4 and 5**). We could particularly observe that short TFP (less than 4 µm) efficiently extend against the flow direction. These short TFP extend straight out (**Fig. 4A**), then either inefficiently retract without generating displacements of the cell body (**Fig. 4B**), or successfully remain attached during retraction thereby powering upstream migration (**Fig. 4C**). In contrast, hydrodynamic forces bend longer TFP along the flow direction (**Fig. 4A**), thus reducing the amplitude of their fluctuations, and reducing the probability of their encounter with the surface.

**Figure 4:**
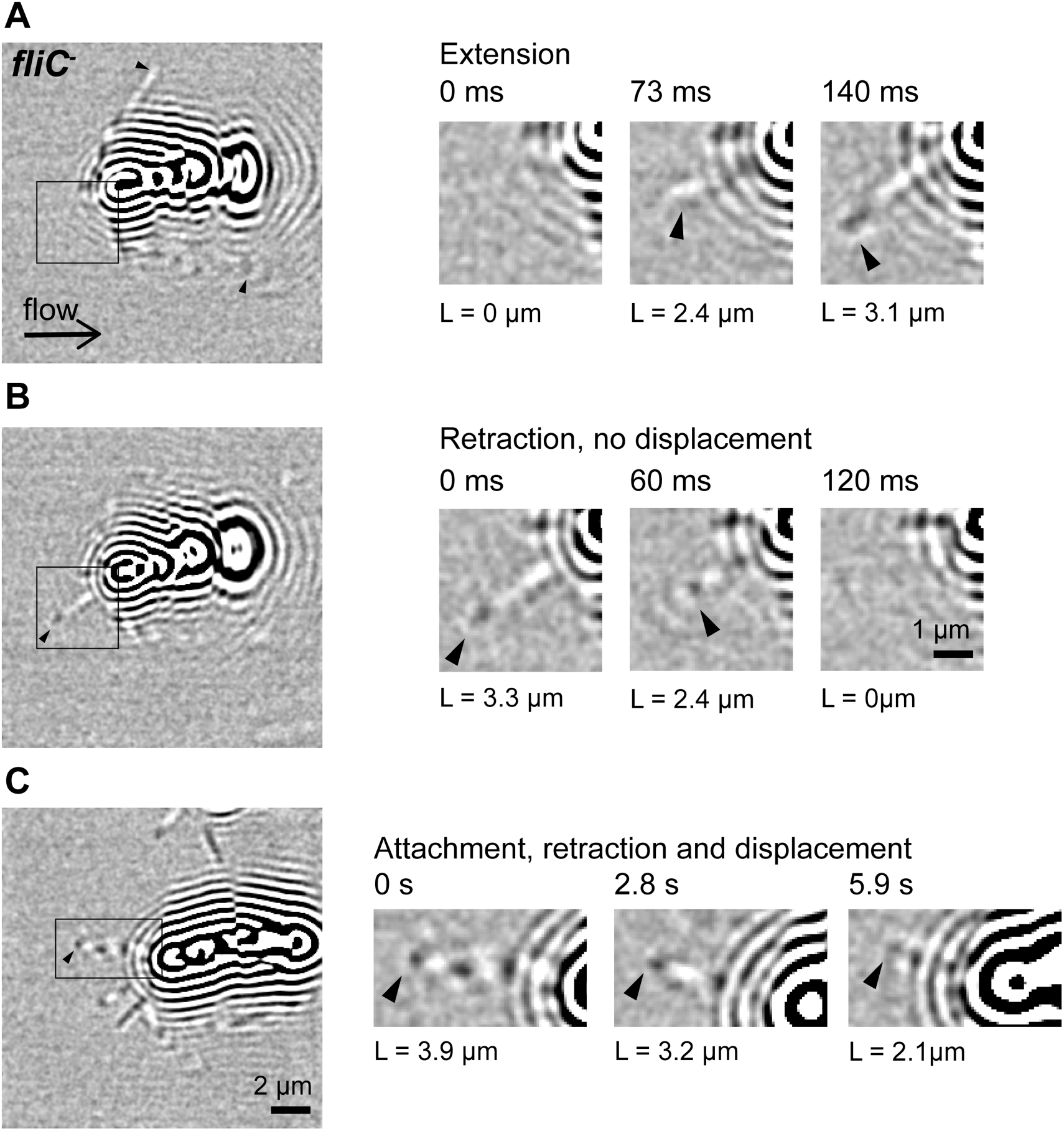
iSCAT reveals that short TFP promote migration against flow. We imaged *P. aeruginosa* migrating against the direction of the flow within a microchannel at 150 fps. (A) Unattached, long TFP are bent by the flow while shorter ones extend straight out against the direction of flow. The image sequence on the right corresponds to the location indicated by a square in the left picture. In this sequence, a pilus extends 3.1 µm within 140 ms. The arrowhead indicates the tip of the pilus. (B) A 3.3 µm-long pilus initially extended is undergoing retraction within 120 ms without generating cell displacement. (C) An attached pilus is retracting, pulling the cell body 1.8 µm forward in 5.9 s (at 0.3 µm.s^-1^ velocity), while its tip remains attached at a stationary location, enabling upstream motility.

### TFP attachment and retraction

Using high-speed iSCAT visualization, we aimed at observing the transitions between extending, attaching and retracting TFP (**Movie 6 and 7**). Here, TFP are fluctuating as they extend from the cell body. Occasionally, a pilus tip would appear as a high contrast, stationary spot while the remainder of the fiber fluctuates, thus indicating tip attachment without retraction. The same fiber subsequently appears as a straight stationary line with uniform contrast along its length indicating retraction (**Fig. 5A**). In our visualizations of *fliC*^-^, all tip attachment events were followed by retraction (**Fig. 5B**). In contrast, we could not observe retraction of unattached TFP, although we have limited sensitivity on detecting filaments that are fluctuating far from the coverslip. The delay between tip attachment and retraction was about 293 ms ± 70 ms (standard error about the mean, *N* = 7), indicating immediate retraction after attachment at the time-resolution of our experiment. This suggests that *P. aeruginosa* coordinates TFP retraction in response to a signal generated upon tip contact with the surface.

**Figure 5:**
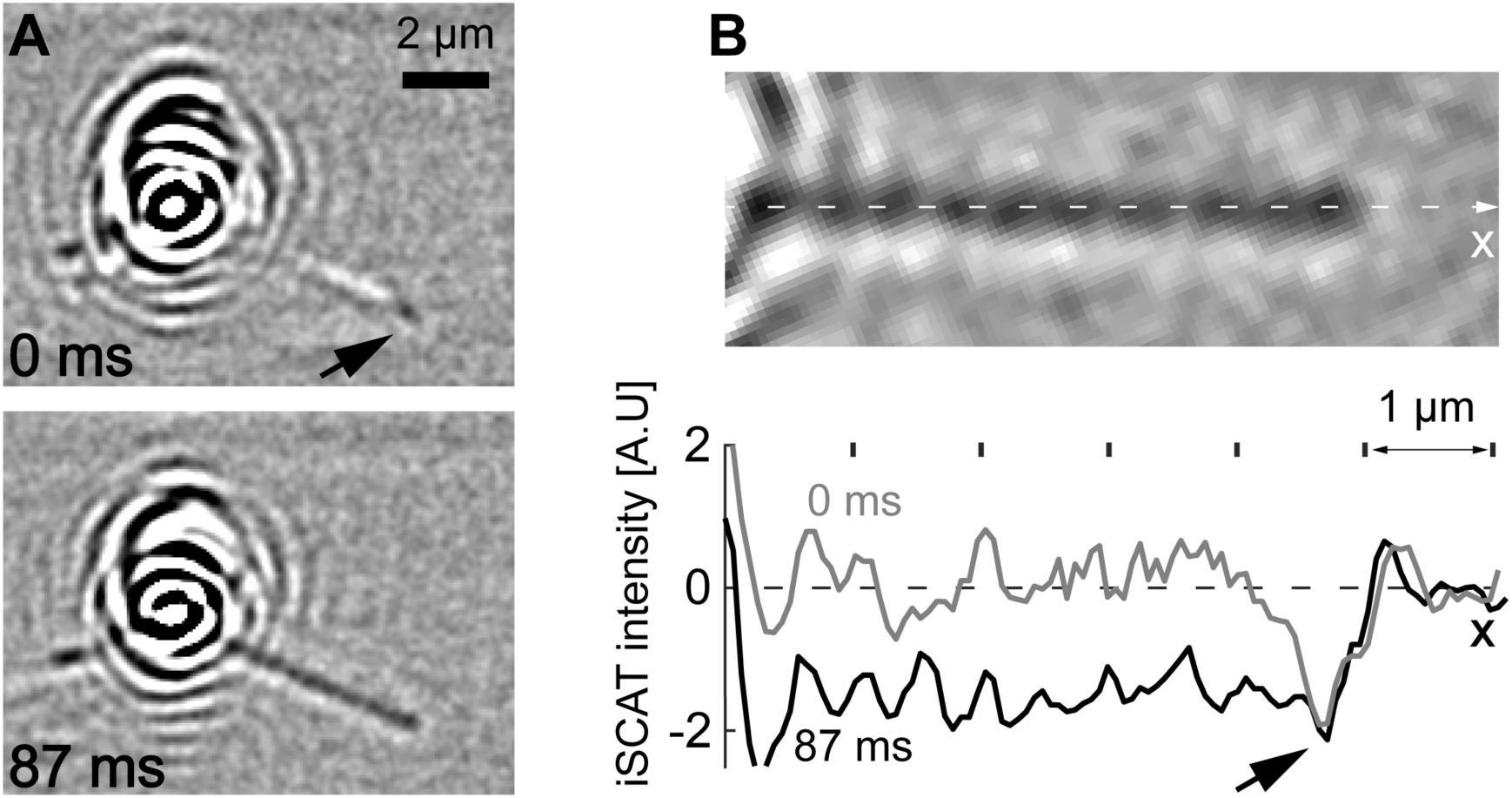
iSCAT reveals a rapid transition between TFP attachment and retraction. (A) The tip of a TFP appears as a stationary dark iSCAT signal (arrow on the image at 0 ms). The remainder of the fiber is fluctuating in space and time with higher intensity values. The fiber subsequently undergoes a transition into an all dark, stationary signal where the TFP is straight, indicating retraction and orientation parallel to the surface (bottom). This transition occurs here in less than 87 ms. (B) Upon initial attachment, the iSCAT intensity in the mid-section of the fiber is slightly positive along the fiber and negative at the tip (arrow on the intensity graph). The signal abruptly transitions to a negative contrast value 87 ms after attachment. Signal from the cell body generates the residual oscillatory contrast away from the cell body.

## Conclusion

By adapting iSCAT to live-cell imaging, we were able to characterize cellular and biophysical functions of TFP without the use of labels. We demonstrated the ability to monitor TFP extension, attachment, retraction, length and orientation at various timescales. Because iSCAT is a label-free technique, we can monitor pili activity without perturbing cellular states thus avoiding conflict with the functionality of the motorized machinery. This feature is key to understand how *P. aeruginosa* and other piliated species regulate twitching motility, activate mechanotransduction and other surface-specific phenotypes using TFP. Finally, since iSCAT is sensitive to all scattering components in the extracellular space, it has the potential to reveal the dynamics of other extracellular machinery in bacteria or in other microorganisms.

With these visualizations, we were able to test functional aspects of the TFP machinery. In particular, we could show that short TFP enable upstream migration of twitching *P. aeruginosa* cells. Also, by measuring extension, attachment and retraction at high speed, we were able to observe a rapid transition between TFP tip attachment and retraction. TFP had been proposed to change conformation during retraction, potentially mechanically activating the Chp chemosensory system^6^. Our new observations suggest that *P. aeruginosa* TFP may in fact sense contact with the surface prior retraction.

## Acknowledgements

The authors would like to thank Joanna Andrecka for valuable discussions on iSCAT, Joanne Engel and Yuki Inclan for strains and plasmids, Zainebe Al-Mayyah for help with generating mutants. LT and AP thank the Swiss National Foundation for funding this work through the Projects grant 31003A_169377.

## Material and Methods

### iSCAT channel

This setup has been adapted from the protocol published by Ortega and Kukura^23^. A 700 mW, 638 nm wavelength with 4 x 5 mm beam diameter and 8 nm bandwidth diode laser (LDM-638-700-C, Lasertack) was used for our iSCAT channel. The beam was spatially filtered through a 50 µm pinhole and collimated with a 4*f* lens system to obtain an output intensity of 99 mW. The center of the collimated beam was aligned into two perpendicular acousto-optic deflectors (AODs), (DTSXY-400-660, AA Opto-Electronic). An iris helped preserve the (1,1) diffraction order while clipping the others. The acquisition software, through a frame grabber (PCIe-1433, National Instruments), triggered the scanning to sync it with the acquisition rate of the camera. The trigger signal activated a burst mode on two function generators (DG1022A, Rigol) that generated a modulation consisting of a ramp function (offset of 1.4 V, amplitude of 1.6 V) with a slight difference between the horizontal and vertical scanning frequencies (79 kHz and 83 kHz respectively. Light intensity was adjusted with a translating linear neutral density filter (NDL-25C-4, Thorlabs). The scanned beam was sent to the back focal plane of an oil immersion objective (PLAN APO 60x 1.42, Olympus) with a 4*f* telescope system inducing a 1/3 beam size shrinkage. In order to split the illumination path from the imaging path, the beam was reflected by a polarizing beam splitter (PBS251, Thorlabs) and a 600 nm short pass dichroic mirror (Part No: 69204, Edmunds optic). A 90° shift in the polarization was obtained through a quarter wave plate (AQWP05M-600, Thorlabs), placed right before the objective back aperture. This shift allows the imaging light to be transmitted by the polarizing beam splitter and to separate it from the illumination light. The signal is then sent to a high dynamic range, high frame rate CMOS camera (pixel size: 31.8 nm, MV1-D1024E-160-CL, PhotonFocus) through a 1000 mm focal length achromatic lens and a 500 nm long pass filter (Part No: 62983, Edmunds optic) giving a total magnification of the setup of 333x. In order to limit iSCAT illumination and reduce photoxicity, we placed a mechanical shutter (SH05/M, Thorlabs) in the illumination path before the PBS. The shutter opened half a second before recording a movie and closed right after acquisition ended. This setup was coupled with a brightfield channel by installing a white LED (WFA1010, Thorlabs) above of the objective. The light captured by the objective was transmitted through the iSCAT 600 nm and an 805 nm short pass dichroic mirrors (used for the autofocus system explained below). It was then focused onto a CMOS camera (CM3-U3-31S4M, PointGrey) by a 400 mm focal length achromatic lens and a 520 nm band pass filter (FBH520-10, Thorlabs) to reject iSCAT and autofocus light, thus achieving a total magnification of 166x.

### Autofocus system

Stability in the focus plane is a critical factor affecting iSCAT image quality. To control this, we built an autofocus system using an 850 nm, 3.5 mW laser (CPS850, Thorlabs). The beam was aligned into an optical fiber and the output Gaussian beam (500 µW) was enlarged and collimated in order to flood the entire objective back aperture. The beam was then sent to the objective by a 50/50 beam splitter (BSW26, Thorlabs) and a 805 nm short pass dichroic mirror (DMSP805, Thorlabs) and was transmitted by the 600 nm short pass dichroic of the iSCAT channel. This dichroic (Part No: 69204, Edmunds optic) has a 20% transmittance for wavelength above 850 nm which yielded sufficient signal. The light at the outer edge of the objective back aperture is totally reflected by the glass water interface resulting in a ring shape pattern. This ring-shaped beam is reflected by the 805 nm short pass dichroic and is partially transmitted by the 50/50 beam splitter. It is then imaged by a CMOS camera (DCC1545M, Thorlabs) through a 100 mm focal length plano-convex lens and an 850 nm band pass filter (FBH850-10, Thorlabs). Computing the mean radius of the ring gives a relative measurement of the Z position of the coverslip. This feeds back onto the voltage of a piezo-stage (KPZNFL5/M, Thorlabs) to maintain ring radius and thus Z position.

### Image acquisition and processing

Images were acquired using LabView and a frame grabber (PCIe-1433, National Instruments) through a CameraLink port. The raw images had to be corrected for the contribution of the constant back reflection from the coverslip in order to reveal the interferometric component of the signal. This was achieved by generating a median image from the whole stack of images from a movie of interest. This median image was then used as a pseudo-flatfield: each frame in the movie is divided by the median image. To improve visualization, and remove fringes generated by AOD scanning, we applied a band pass filter dampening the contributions of small and large structure (smaller than 1 pixel and larger than 13 pixel) with the FFT plugin of imageJ. To further normalize the images, each frame was divided by its mean pixel gray value.

### Strains and sample preparation

Strains and plasmids used in this work were described previously^19^. Double deletion mutants *pilH*^-^ *fliC*^-^ and *pilTU*^-^ *fliC*^-^ were generated by conjugation of *pilH*^-^ and *pilTU*^-^ mutants with the plasmid pJB215 using a standard mating protocol^24^.

Single colonies of the specified bacterial strains were grown in LB medium overnight. The culture was diluted 1:200 and grown to mid-exponential phase before visualization. The cells were loaded into 500 µm wide, 90 µm deep PDMS microchannels fabricated using standard photolithography methods^25^. The chip was first loaded with plain LB medium, after proper mounting of the tubing and syringes, the exhaust tubes were dipped in the bacterial culture. After mounting the chip on the microscope and connecting the syringes to a syringe pump, we evacuated excess air in the exhaust tubes prior to cell loading. The bacteria were then loaded by aspiration of the culture medium until cells were visible in the microchannel under brightfield. Cells were imaged either at no flow conditions or under flow conditions (from 0.1 µl/min or 10 µl/min).

---

**Movie 1: iSCAT visualization of the retraction motors mutant *pilTU*^-^**.

Real time observation of the *pilTU*^-^ mutant shown in **Fig. 1E**. This mutant display many surface type IV pili that extend and occasional attach on the coverslip but never retract. The acquisition rate was 200 frames per second (fps), binned in time twice thus reducing frame rate to 100 fps. The movie is under-sampled and played in real time at 20 fps. The width of the field of view is 16.3 µm.

**Movie 2: iSCAT visualization of the response regulator mutant *pilH*^-^**.

Visualization of two cells of the *pilH*^-^ mutant shown in **Fig. 3** The acquisition rate was 5 fps, binned in time 2x thus reducing the frame rate to 2.5 fps. The movie is accelerated 10 times (playing at 25 fps). The width of the field of view is 23.5 µm.

**Movie 3: iSCAT visualization of TFP extension and retraction in flow**.

High frame rate visualization (300 fps acquisition, 2x bin resulting in 150 fps) of pili extending and retracting of the cell shown in **Fig. 4** in flow (flow rate 10 µl.s^-1^). The movie is slowed 30x to better visualize extension and retraction (playing at 5 fps). The width of the field of view is 23.5 µm.

**Movie 4: iSCAT visualization of TFP extension and retraction in flow**.

Same as **Movie 3** played at real speed (150 fps).

**Movie 5: iSCAT visualization of TFP extension and retraction generating upstream twitching in flow**

Same imaging parameters as **Movie 4**.

**Movie 6: iSCAT visualization of extension, attachment and retraction of a TFP**.

High frame rate visualization (300 fps acquisition, 2x bin resulting in 150 fps) of pili extending, attaching and retracting of the cell shown in **Fig. 5**. The movie is slowed 30x (5 fps) to better visualize the time delay between attachment and retraction until the pili is under tension. The width of the field of view is 23.5 µm.

**Movie 7: iSCAT visualization of extension, attachment and retraction of a TFP**.

Same as for **Movie 6** but played at real speed (150 fps).

